# Unifying microorganisms and macrograzers in rocky shore ecological networks

**DOI:** 10.1101/2023.10.06.561312

**Authors:** Clara Arboleda-Baena, Claudia Belén Pareja, Javiera Poblete, Eric Berlow, Hugo Sarmento, Ramiro Logares, Rodrigo De la Iglesia, Sergio A. Navarrete

## Abstract

Over the past decades, our understanding of the vital role microbes play in ecosystem processes has greatly expanded. However, we still have limited knowledge about how microbial communities interact with larger organisms. Many existing representations of microbial interactions are based on co-occurrence patterns, which do not provide clear insights into trophic or non-trophic relationships. In this study, we untangled trophic and non-trophic interactions between macroscopic and microscopic organisms on a marine rocky shore. Five abundant mollusk grazers were selected, and their consumptive (grazing) and non-consumptive (grazer pedal mucus) interactions with bacteria in biofilms were measured using 16S rRNA amplicon sequencing. While no significant effects on a commonly used measure of biofilm grazing (Chlorophyll-a concentration) were observed, detailed image analysis revealed that all grazers had a detrimental impact on biofilm cover. Moreover, different grazers exhibited distinct effects on various bacterial groups. Some groups, such as Rhodobacteraceae, Saprospiraceae, Flavobacteriaceae, and Halieaceae, experienced positive effects from specific grazers, while others, like Rhizobiaceae, Rhodobacteraceae, and Flavobacteriaceae were negatively affected by certain grazers. This study presents the first attempt to construct an interaction network between macroorganisms and bacteria. It demonstrates that the strength of trophic and non-trophic interactions varies significantly depending on the mollusk grazer or bacterial group involved. Notably, certain bacterial groups exhibited a generalized response, while others showed specialized responses to specific macroorganisms in trophic or non-trophic interactions. Overall, this work highlights the potential for integrating microbes into ecological networks, providing valuable insights and methodologies for quantifying interactions across Domains. This research complements the previous ecological network, showing that mollusk grazers interact not only trophically but also non-trophically with epilithic biofilms. It identifies three drivers affecting microbial community assembly, crucial for understanding macro-microorganism dynamics in intertidal systems.

## INTRODUCTION

Organisms in ecosystems engage in complex interactions with multiple species, creating intricate networks where changes in one species can affect seemingly unrelated species in unpredictable ways (Yodzis 2001). Beyond trophic interactions, where organisms obtain energy by consuming biomass (Lindeman 1942), there are numerous non-trophic interactions with diverse effects, including positive ones like commensalism, facilitation, and mutualism, as well as negative ones like inhibition and interference (Kéfi et al. 2012).

Network representations, where co-occurring species are nodes and a single type or multiple types of interactions (multiplex networks) as vertices or links, offer a framework to study the structure and dynamics of complex ecological networks (Bascompte 2010). Substantial progress has been made in recent years, enabling realistic visualization and characterization of these networks (Sander et al. 2015, 2017, Kéfi et al. 2016) and modeling their dynamics (Rebolledo et al. 2019, Valdovinos 2019, Ryser et al. 2021).

Coastal marine ecosystems have provided some of the most detailed ecological networks due to extensive experimental research on trophic and non-trophic interactions (Kéfi et al. 2015, 2016, Sander et al. 2015, 2017). However, a common weakness in well-documented marine and terrestrial food webs (Link 2002, Lafferty et al. 2008, Digel et al. 2014), as well as multiplex ecological networks (Hale et al. 2020, Costa et al. 2020), is the neglect or misrepresentation of co-occurring microorganisms that interact with macroorganisms (Sechi et al. 2015).

Microorganisms are the most abundant and diverse organisms on Earth (Locey and Lennon 2016). They are essential for regulating biogeochemical cycles (Gasol and Kirchman 2018), providing materials and energy to higher trophic levels (Castenholz 1961, Thompson et al. 2000). While our understanding of microorganisms at the ecosystem level is well-established, we remain profoundly ignorant about their interactions with macroorganisms at the community level (Bjorbækmo et al. 2020). Despite extensive research on microbiomes within macroscopic organisms, we still treat the microbial world as a separate and non-interacting black box. Neglecting the interactions between macroscopic and microscopic communities prevents us from capturing the dynamics of ecological networks realistically and understanding the functions, persistence, and responses of complex ecological systems (Koskella et al. 2017).

To approach this problem, we used as a study model one of the most resolved ecological networks, the intertidal rocky shore of central Chile (Kéfi et al. 2015, 2016). Our study aimed to uncover the trophic and non-trophic connections between the prevalent intertidal mollusk grazers and the bacterial components of the co-occurring epilithic biofilms (periphyton) (**Fig. 1**) (Arboleda-Baena et al. 2021).

**Fig. 1.**
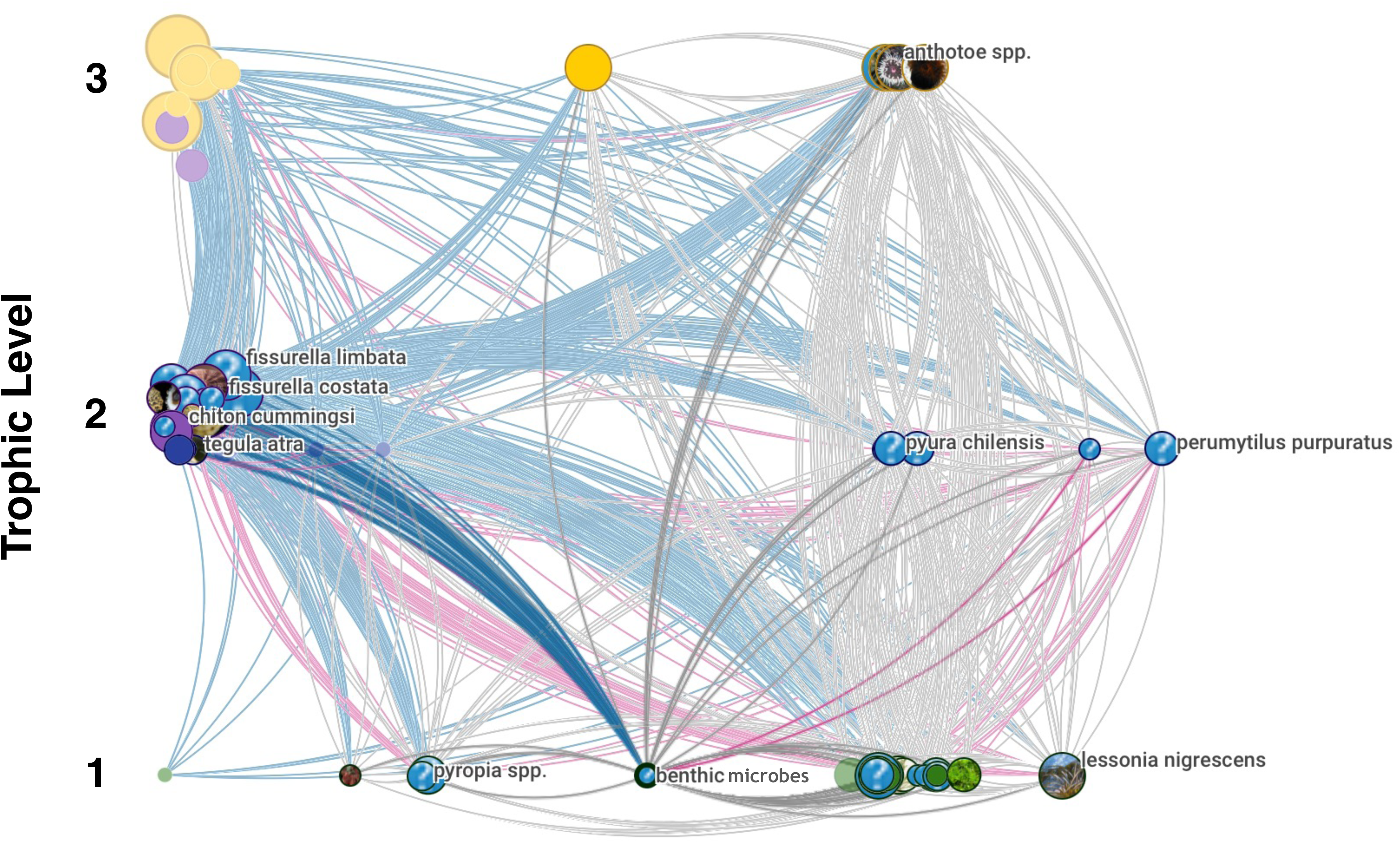
Ecological network of known species interactions in the Chilean marine rocky shore ecosystem. Nodes represent species, and the links depict three types of interactions: 1) Feeding (blue), 2) Competition (grey), and 3) Facilitation (pink). Herbivore-diatom interactions are highlighted. Image adapted from https://map.openmappr.org/chile-marine-intertidal-network/, Vibrant Data Labs.

In trophic interactions, epilithic biofilms are essential for the primary productivity of coastal ecosystems (Bustamante et al. 1995, Thompson et al. 2000 p. 202, Jenkins et al. 2001), and on many shores, they are the primary food resource for many benthic grazers (Castenholz 1961, Thompson et al. 2000, Christofoletti et al. 2011). The molluscan grazers feed by scraping rock surfaces, removing algae, invertebrate settlers, and microorganisms to varying extents depending on the species and their ecological and morphological traits (Underwood 1984, Santelices et al. 1986, Jenkins et al. 2001, Christofoletti et al. 2011, Aguilera and Navarrete 2012). Meanwhile, non-trophic interactions involving grazers and epilithic biofilms have been shown to have both positive and negative effects, mediated by the grazers’ pedal mucus (Davies et al. 1990, Arboleda-Baena et al. 2022).

But how many of the microbial ‘species’ that comprise the biofilms are negatively or positively affected by trophic consumption of a given grazer species? Which type of microbial organisms are more susceptible and therefore more significantly affected? And at which trophic level within the microbial community, is yet unknown for this and all marine ecosystems? Disentangling this web of interactions and quantifying the strength of trophic and non-trophic interactions requires experimental approaches (Paine 1992, Wootton 1997, Berlow et al. 1999, Aguilera and Navarrete 2012). This will allow us to start developing predictive models of dynamic ecological networks (Laska and Wootton 1998, Berlow et al. 1999).

Here we use experiments to quantify the interaction strengths, both trophic (consumptive) and non-trophic (mediated through pedal mucus), among five model species from distantly related mollusk taxa and the bacterial components of the biofilm community. We test whether grazers exert stronger effects on the abundance of epilithic biofilms (cover and Chlorophyll-a concentration) and if they have stronger or more widespread effects on the dominant bacterial taxa. Furthermore, we investigated whether trophic interactions (TI) between grazers and epilithic biofilms have a greater impact on the bacterial community compared to non-trophic interactions (NTI). Additionally, this study explores variations in trophic and non-trophic interactions within an intertidal macrograzer assemblage, focusing on their influence on the abundance and diversity metrics of epilithic biofilms.

## MATERIALS & METHODS

### Study site and mollusk grazer assemblage

The study was conducted at the Estación Costera de Investigaciones Marinas (ECIM) of Pontificia Universidad Católica de Chile, located in Las Cruces, Chile (33° 30’ S, 71° 38’ W). We chose five of the most abundant grazer species in terms of total biomass (Arboleda-Baena et al. 2022), one Polyplacophoran, the chiton *Chiton granosus* (Frembly 1828) Family Chitonidae, and four Gastropods, the Littorinid *Echinolittorina peruviana* (Lamarck 1822) Family Littorinidae, the keyhole limpet *Fissurella crassa* (Lamarck 1822) Family Fissurellidae, the scurrinid limpet *Scurria araucana* (d’Orbigny 1839) Family Lottiidae, and the pulmonate limpet *Siphonaria lessonii* (Blainville 1827) Family Siphonariidae (Espoz et al. 2004, Aguilera et al. 2013). These omnivores scrap rock surfaces, primarily removing periphyton (epilithic biofilm), ephemeral algae, and newly established invertebrates (Camus 2008, Aguilera and Navarrete 2012). We measured their body size (**Appendix S1: Table S1)** and classified them by their radula, from greater to lesser excavation capacity, in the following order (Radula type in parenthesis): *C. granosus* (Steroglossa)*, S. araucana* (Docoglossa)*, S. lessonii* (Taenioglossa)*, E. peruviana* (Taenioglossa), and *F. crassa* (Rhipidoglossa) (Steneck and Watling 1982, Reid 2002). Field study approval number ID Protocol: 170829006, by the Comité Institucional de Seguridad en Investigación of the Pontificia Universidad Católica de Chile.

### Grazer and epilithic biofilm sampling

First, field rocks were collected and cut into 3 x 8 x 2 cm coupons using a COCH Bridge saw machine to prevent overheating and mineral modification. The coupons were then cleaned with deionized water, dried, and kept at room temperature until introduced into the experimental aquaria. To establish epilithic biofilm communities, the rock coupons were placed in aquaria with circulating seawater sourced from the same location and provided with continuous aeration between September and October 2018.

Second, we collected fifty individuals of each grazer species from gently sloping wave-exposed platforms near ECIM. The collection took place during nocturnal low tides to avoid foot damage (Aguilera and Navarrete 2012). The collected animals were transported in coolers to the laboratory and then, to reduce animal stress, acclimatized for a week in separate aquaria with circulating seawater and constant aeration. During this period, the grazers were exclusively fed with epilithic biofilm provided in the same manner. Then, to minimize fecal contamination before the experiment, the grazers underwent a two-day cleaning period in an aquarium with constant aeration and 0.2 µm filtered seawater (from the same location). During this period, the animals starved, reduced fecal production, and avoided adverse locomotory and metabolic effects (Calow 1974). On the third day, reduced motility was observed, indicating the effective prevention of locomotory and metabolic disturbances during the cleaning period (data not shown). The filtered water was replaced every 2-6 hours to minimize ammonium concentration and prevent biofilm formation associated with animals’ feces. This cleaning period also helped remove incidental microorganisms on the animal foot that do not maintain populations in the pedal mucus. The motility and behavior of the animals were monitored throughout the acclimation period. Seawater temperature was maintained at 13° C +/− 2°C, which was the average SST during the experiments.

### Grazer effects on integrated measures of epilithic biofilm abundance

To quantify grazing effects on total Chlorophyll-*a* content and cover of epilithic biofilm, we conducted a replicated laboratory experiment at ECIM described in **Appendix S1: Figure S1.**

### Trophic and non-Trophic Grazer effects on epilithic biofilm community composition

To quantify the trophic (grazing) and non-trophic (pedal mucus) interactions, a second experiment was conducted with eleven different treatments (a - k). Six treatments were implemented to examine the trophic effects (a - f), with 13 replicates each: a) Control: epilithic biofilm rock control without grazers. One rock coupon with epilithic biofilm plus one individual of either b) *C. granosus*, c) *E. peruviana*, d) *F. crassa*, e) *S. araucana*, f) *Siphonaria lessonii.* Simultaneously we assessed the effect of pedal mucus by including five treatments (g - k) with 10 replicates each. In this case, rock coupons with epilithic biofilm were surrounded by a sterile transparent plastic mesh cage that impeded grazing on the surface. One individual of either g) *C. granosus*, h) *E. peruviana*, i) *F. crassa*, j) *S. araucana*, or k) *S. lessonii* was included in each experimental aquaria. In these treatments, grazers moved on top of the protective mesh, and their pedal mucus was in contact with the filtered marine water on top of the rock coupons with epilithic biofilms. The pedal mucus microbiota entered in contact with the epilithic biofilms, but the grazers could not move on top of the rock to graze it. See **Appendix S1: Fig. S1.**

In this manner, we can examine: 1) Total grazer effects (T), in which treatments b-f, where grazers feed in biofilm and pedal mucus is in contact with the rock’s surface, are compared against controls without grazers; 2) non-Trophic interactions (NTI), in which treatments with only pedal mucus (g-k) are compared against the respective controls, and 3) Trophic interaction (TI), which is obtained by subtracting the respective pedal mucus effect from the total effect. Treatments were randomly assigned to the 128 experimental units. Every three hours, the temperature was checked, and feces were carefully removed. The experiment lasted 24 hours, and then the animals were carefully removed. Longer exposure was not possible because some grazers had already significantly reduced biofilm cover after 24 hours and because we wanted to focus on the direct impacts of grazing on the resulting biofilm community. Thus, to analyze the effect of grazers on bacterial communities reassemblage, rocks were marked with graphite on the grazed zone, separating that zone from the rest of the epilithic biofilm community that remained on the rock. Rocks were transferred to new aquaria containing circulating seawater from the same location, along with constant aeration, for 10 days. This specific period was determined based on the observation of complete biofilm recovery on the rocks (data not shown). Then, after 10 days of recovery, 10 replicates of each treatment were sampled with a sterile scalpel from control rocks, grazed areas of grazing treatment and pedal mucus treatment, placed on 0.22 µm pore filters of hydrophilic polyether sulfone (Merck), and preserved in liquid nitrogen at −196°C for a subsequent DNA extraction and 16S rRNA-gene sequencing. During the molecular analyses, we lost samples due to a poor-quality Illumina sequencing run (see **Appendix S1: Table S2**).

Three replicates from each of the trophic interaction (grazing) treatments (treatments labeled as a-f) were subjected to analysis for total Chlorophyll-a concentration after a 10-day recovery period, following the identical protocol described above. The Chlorophyll-a concentrations obtained from the grazed area of the rock coupon were standardized by area. Subsequently, to assess the statistical significance, Welch’s ANOVA with six levels as a fixed factor was conducted after confirming the absence of homoscedasticity.

### DNA extraction and 16S rRNA-gene sequencing

DNA extraction from filters was conducted with the Phenol-Chloroform method (Fuhrman et al. 1988). DNA concentration was measured with the Qubit HS dsDNA Assay kit in a Qubit 2.0 Fluorometer (Life Technologies, Carlsbad, CA, USA) according to manufacture protocols. The V4-V5 region of the 16S rRNA gene was amplified with the primers 515FB: GTGYCAGCMGCCGCGGTAA and 926R: CCGYCAATTYMTTTRAGTTT (Parada et al. 2016). Amplicons were sequenced in a MiSeq Illumina platform (2×300 bp). Both PCR and sequencing were done at the Dalhousie University CGEB-IMR (http://cgeb-imr.ca/). Sequence data will be deposited in the European Nucleotide Archive (ENA) database should the manuscript be accepted for publication.

### Epilithic biofilm community analyses

Amplicon reads were analyzed using the DADA2 pipeline (Callahan et al. 2016) to characterize Amplicon Sequence Variants (ASVs) (Callahan et al. 2017). Rarefaction curves were generated with a fixed sampling effort of 6,215 reads per sample, due to the size of the smallest dataset from one replicate of the epilithic biofilm control (**Appendix S1: Fig. S2)**.

To examine the microbial beta diversity after grazing (trophic interaction, TI) or the grazer pedal mucus effect (non-trophic interaction, NTI) of the five most abundant grazer species, we used non-metric multidimensional scaling (NMDS) ordination, based on Bray-Curtis dissimilarities. To compare the composition among treatments we conducted a permutational analysis of variance (PERMANOVA) (Anderson and Walsh 2013). To determine which treatment differed from others, after a significant Permanova, we conducted pairwise post hoc tests with False Discovery Rate (FDR) correction for multiple comparisons (Benjamini and Hochberg (1995). To compare richness and diversity among treatments, we conducted a separate one-way ANOVA or Kruskal-Wallis, respectively, after corroborating approximate homoscedasticity in both cases and large deviations from normality in the case of diversity and considered treatment (grazer species and control) as a fixed factor. We used Tukeýs or Dunn post hoc test to establish the pattern of differences.

All graphics and statistical analyses were carried out in R with RStudio interface (Racine 2012). Most community analyses were carried out using packages vegan v2.5-6 (Oksanen et al. 2013) and phyloseq v1.30.0 (McMurdie and Holmes 2013)

### Interaction strength calculation between grazers and microbial populations

Quantification of grazer per capita effects on the relative abundance of microbial groups was estimated by the Dynamic Index (DI) (Berlow et al. 1999). Where DI = (LN(N/D))/Yt; N is the relative abundance of ASVs in the treatment where grazers are present (i.e., “grazing treatments); D is the relative abundance of ASVs in the treatment where grazers are absent (i.e., “Control of epilithic biofilm”); Y, the abundance of the predator (i.e., our experiment used an abundance of 1 per replicate); t, time (i.e., our experiment only used one time). This index is recommended for short-term experiments, comparing both positive and negative effects and when the resources exhibit a positive exponential growth (Berlow et al. 1999). Confidence intervals (95%) were obtained through a bootstrapping procedure (Manly 2006). The calculation was performed with ASVs with a number of reads ≥ 0.1%, to eliminate the rare bacteria (Pedrós-Alió 2012). To illustrate patterns in the interaction strengths of grazers on different microbial groups we showed only those links with an absolute DI score of at least 1 for the most abundant microbial ASVs (for those ASVs with a total number of reads > 0.5%) (Pedrós-Alió 2012).

### Bipartite networks analyses

We constructed bipartite networks of positive and negative interactions between macrograzers and microbial ASVs with interaction strength values higher than 1. The network visualization was performed with the software Cystoscape (https://cytoscape.org). Specialization indices of bipartite networks (Blüthgen et al. 2006, 2007) were calculated at the network level (*H_2_′*) and at bacterial group level (*d′*) using H2fun and dfun functions from bipartite R package (Dormann et al. 2008) in Software R (http://www.r-project.org) after multiplying values by 1000 and rounding them to integers, as calculations of specialization indices require the absence of decimals. The index d’ quantifies the extent to which species within one partition (e.g., bacteria) interact with a subset of species in the other partition (e.g., grazers) more than expected by chance. The index *H_2_′* measures nestedness in bipartite networks, a higher degree of nestedness means that species with few interactions tend to interact with subsets of the species with higher interactions. Specialization indices *H_2_′* and *d′* range from 0 for the most generalized to 1 for the most specialized case (Blüthgen et al. 2006).

## RESULTS

### Grazing effects in integrated measures of biofilm community abundance after 24 hours

Although the Chlorophyll-a concentration (mg m-2) was lower under the *F. crassa* treatment and, to a lesser extent, under *S. lessoni* after 24 hours of grazing, there were no statistical differences among the grazer species and the control group (**Fig. 2a**, Welch’s ANOVA, *df* = 5, p-value = 0.22). The per capita effect on the Chlorophyll-a content of epilithic biofilms, as calculated using the Dynamic Index (DI) (**Appendix S1: Fig. S3a**), indicated that both *F. crassa* and *S. lessonii* had a significant impact over epilithic biofilm within a 95% confidence interval. In contrast, all grazer treatments had a significant negative effect on the cover of epilithic biofilms (mean% ± SE) compared to the control group (99.9 ± 0.4) (**Fig. 2c**). However, the chiton *C. granosus* (54 ± 4.8) and the keyhole limpet *F. crassa* (44.2 ± 2.8) had significantly greater effects compared to *E. peruviana* (91.4 ± 1.0), *S. araucana* (85.2 ± 1.5), and *S. lessonii* (83.2 ± 1.9). The cover analysis of the same experiment revealed statistical differences among all the grazer species treatments (**Fig. 2c**, Welch’s ANOVA, *df* = 5, p < 0.0001) (Appendix **S1: Table S2**).

**Fig. 2.**
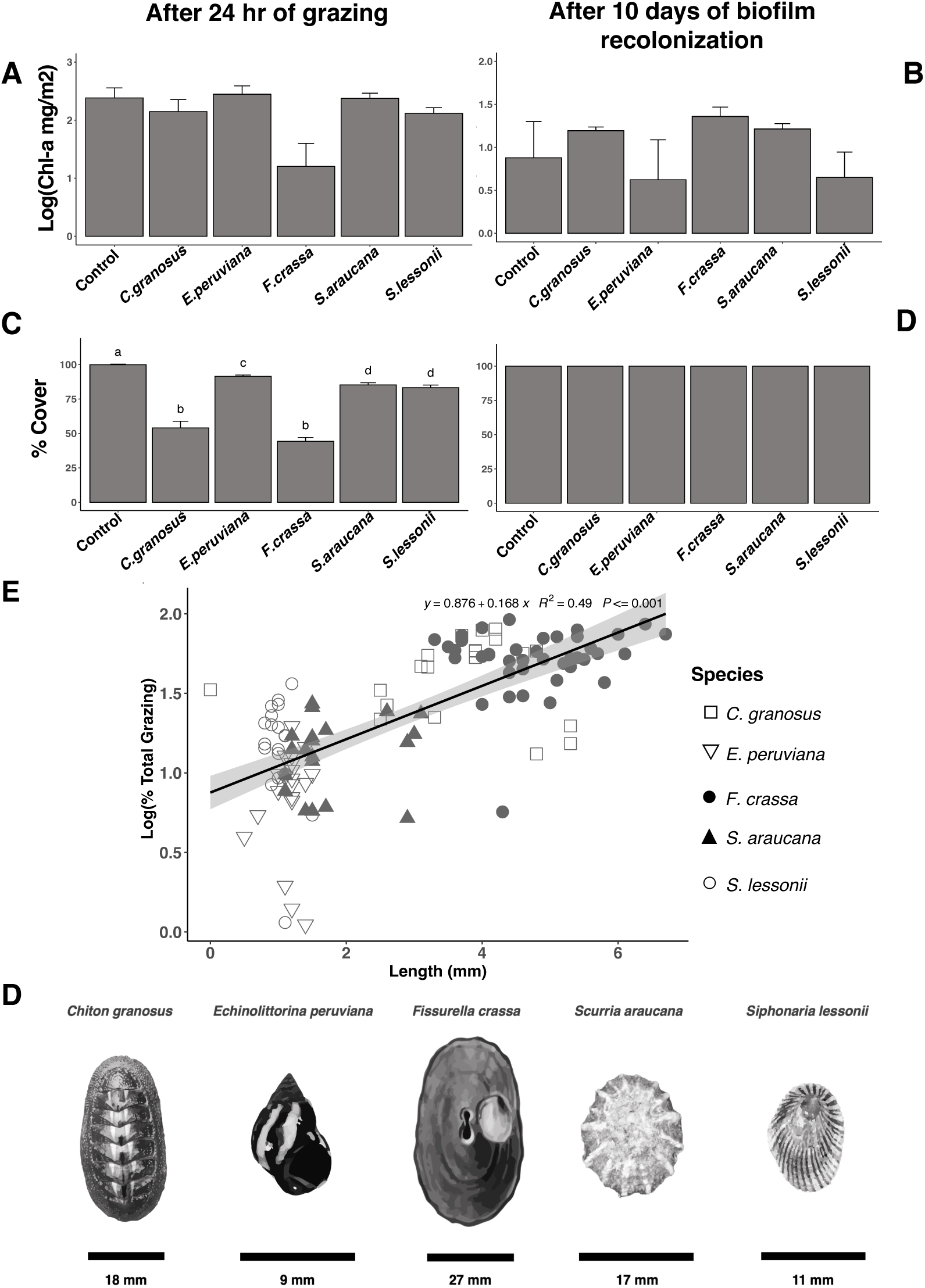
Grazing effect on epilithic biofilm. A) Biofilm Chlorophyll-a content and biofilm cover after 24 hours of grazing, and B) biofilm recolonization over 10 days in the grazed area (mean + SE). Different letters above bars indicate significant differences among treatments (Games-Howell test, experiment-wise error rate = 0.05). E) Linear regression showing the relationship between total grazing percentage and body size (mm) of *Chiton granosus* (□), *Echinolittorina peruviana* (▽), *Fissurella crassa* (●), *Scurria araucana* (▲), and *Siphonaria lessonii* (○). F) Grazers species selected for analysis. Scale bar is provided for each species. Images credits: Ana María Valencia.

The per capita effect, calculated using the Dynamic Index (DI), yielded consistent results, with *C. granosus* and *F. crassa* having the highest impact, followed by *S. araucana* and *S. lessonii*, and finally *E. peruviana* treatments (**Appendix S1: Fig. S3c**). Among the grazers, *F. crassa* (42.9 mm ± 3.9) was by far the largest, closely followed by *C. granosus* (34.3 mm ± 2.5), while *S. araucana* (16.1 mm ± 0.8), *E. peruviana* (10.1 mm ± 0.2), and *S. lessonii* (8.7 mm ± 0.1) were smaller in decreasing order (**Appendix S1: Table S3**). Given the correlation between body size and consumption rates, we compared the relationship between grazing percentage and the size of the grazer assemblage. Our analysis revealed a positive correlation between these two variables (R^2^ = 0.49, p-value < 0.001). Consequently, even with a different type of radula, larger grazers have a greater potential to impact the cover of epilithic biofilms (**Fig. 2e**).

We utilized scanning electron microscopy to investigate changes in the coverage and structure of the epilithic biofilm community. Our observations revealed a significant reduction in biomass in treatments involving *F. crassa* and *C. granosus* compared *to E. peruviana, S. araucana,* and *S. lessonii* (**Appendix S1: Fig. S4**). Furthermore, when comparing the images and remaining microorganisms in the first two treatments, we observed that the Steroglossa radula of *C. granosus* caused the destruction of all cell structures. Conversely, the Rhipidoglossa radula of *F. crassa* did not have the same effect on the remaining cells (**Appendix S1: Fig. S4**).

### Grazing effects in integrated measures of biofilm community abundance after 10 days of epilithic biofilm recolonization

Following the grazing experiment, and a ten-day period of biofilm recolonization, there were no significant differences observed in the Chlorophyll-a content (**Fig. 2b**, Welch’s ANOVA, *df* = 5, p-value = 0.45) or the cover (**Fig. 2d**) of the grazed area among the treatments. This indicates a complete recovery of the epilithic biofilm. The per capita effect, as measured by the Dynamic Index (DI), for both Chlorophyll-a content and cover was either negligible or not statistically significant in all treatment groups (**Appendix S1: Fig. S3b, d**).

Statistical analysis using PERMANOVA revealed significant differences in bacterial community similarity following trophic interactions (TI) (*df* = 4, p-value = 0.028, **Fig. 3a**). However, after applying the FDR post hoc test for correction, the differences were no longer statistically significant (**Appendix S1: Table S4**). On the other hand, the composition and relative abundances of bacterial communities remained unchanged after non-trophic interactions across all five grazer treatments (PERMANOVA, *df* = 4, p-value = 0.081, **Fig. 3b**).

**Fig. 3.**
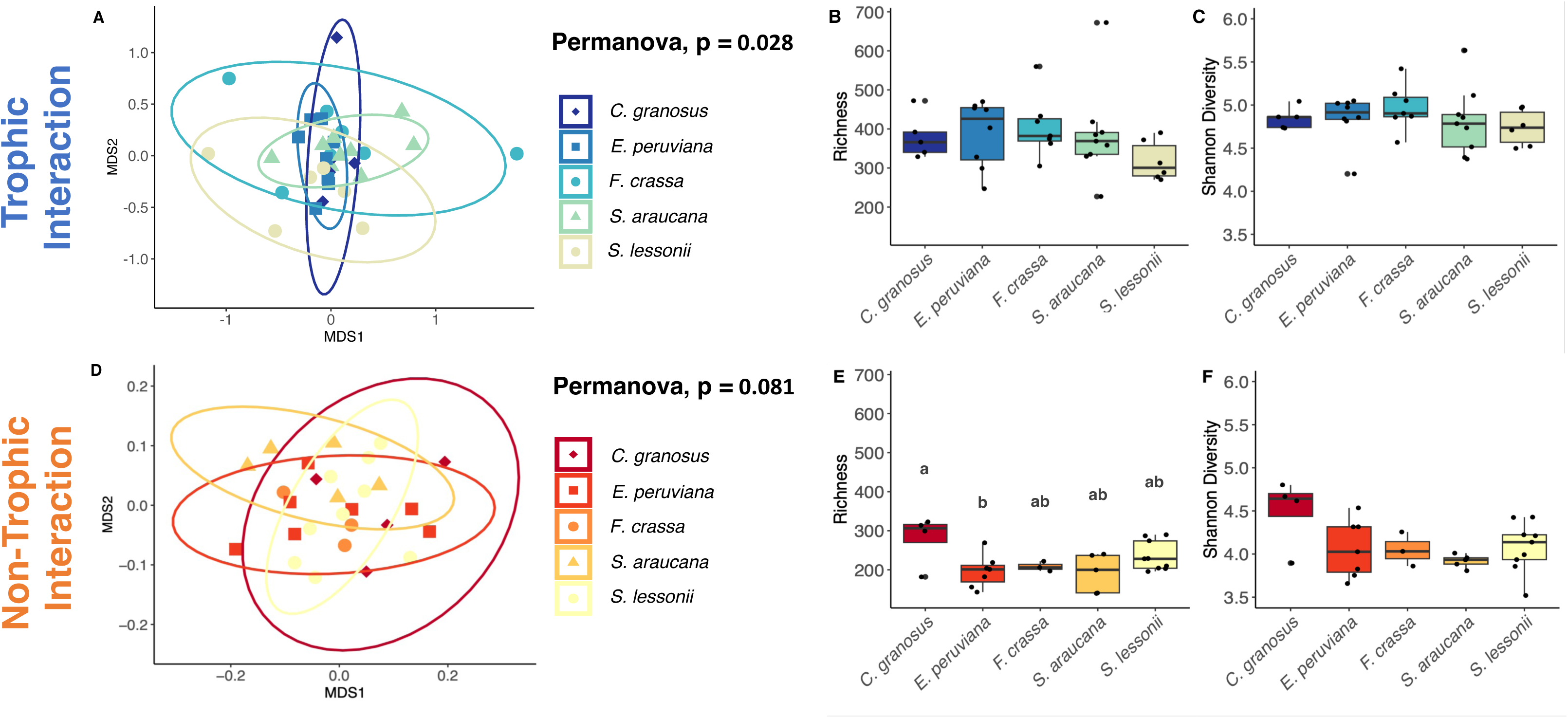
Bacterial community compositional similarity, richness, and diversity after trophic/non-trophic interactions with Molluscan grazers. A, D) Non-metric multidimensional scaling (NMDS) ordination plots based on Bray-Curtis distances. Each grazer species treatment is represented by a specific shape surrounded by a 95% confidence interval ellipse: (♦) *C. granosus*, (▪) *E. peruviana*, (●) *F. crassa*, (▲) *S. araucana*, (●) *S. lessonii*. A) Stress = 0.186, D) Stress = 0.192. B, E) Mean richness, and C, F) mean Shannon diversity index (mean + SE). Different letters above bars indicate significant differences based on the Tukey test, experiment-wise error rate = 0.05.

The comparison of richness and Shannon diversity index of bacterial communities following trophic interactions with the five grazer species did not reveal statistically significant differences (**Fig. 3b, c**; Kruskal-Wallis test, *df* = 4, p-value = 0.288, and ANOVA, *df* = 4, p-value = 0.684, respectively). However, microbial richness exhibited significant variation among the grazer’s treatments after non-trophic interactions (**Fig. 3e**, ANOVA, *df* = 4, p-value = 0.033), while microbial diversity remained similar (**Fig. 3f**, ANOVA, **df** = 4, p-value = 0.079). Specifically, the microbial richness of the *C. granosus* and *E. peruviana* grazing treatment differed significantly (Tukey post hoc test, p-value = 0.04, **Appendix S1: Table S5a**), with *C. granosus* (279 ± 33 richness) showing higher values compared to *E. peruviana* (196 ± 16 richness).

### Interaction strength between grazers and bacterial populations

A bipartite network illustrating positive and negative interaction strengths was constructed (**Fig. 4**). Overall, both trophic and non-trophic interactions exhibited varying magnitudes and directions of individual per capita effects on the abundance of dominant microbial ASVs by *C. granosus, E. peruviana, F. crassa, S. araucana,* and *S. lessonii*, as estimated by the Dynamic Index (DI) (range between −1 < DI < 1) (**Fig. 4** and **Appendix S1: Fig. S5**). The trophic interactions demonstrated higher values of interaction strength, both positive and negative. Following the trophic interaction, 24 ASVs displayed a positive or negative per capita effect by the five studied grazer species. All grazer species exhibited significant impact, with a 95% confidence interval (CI), on ASVs belonging to the Alphaproteobacteria Class, particularly the Family Rhodobacteraceae, except for ASV_60 (*Sedimentatitalea* sp.) (**Fig. 4a** and **Appendix S1: Fig. S5a**).

**Fig. 4.**
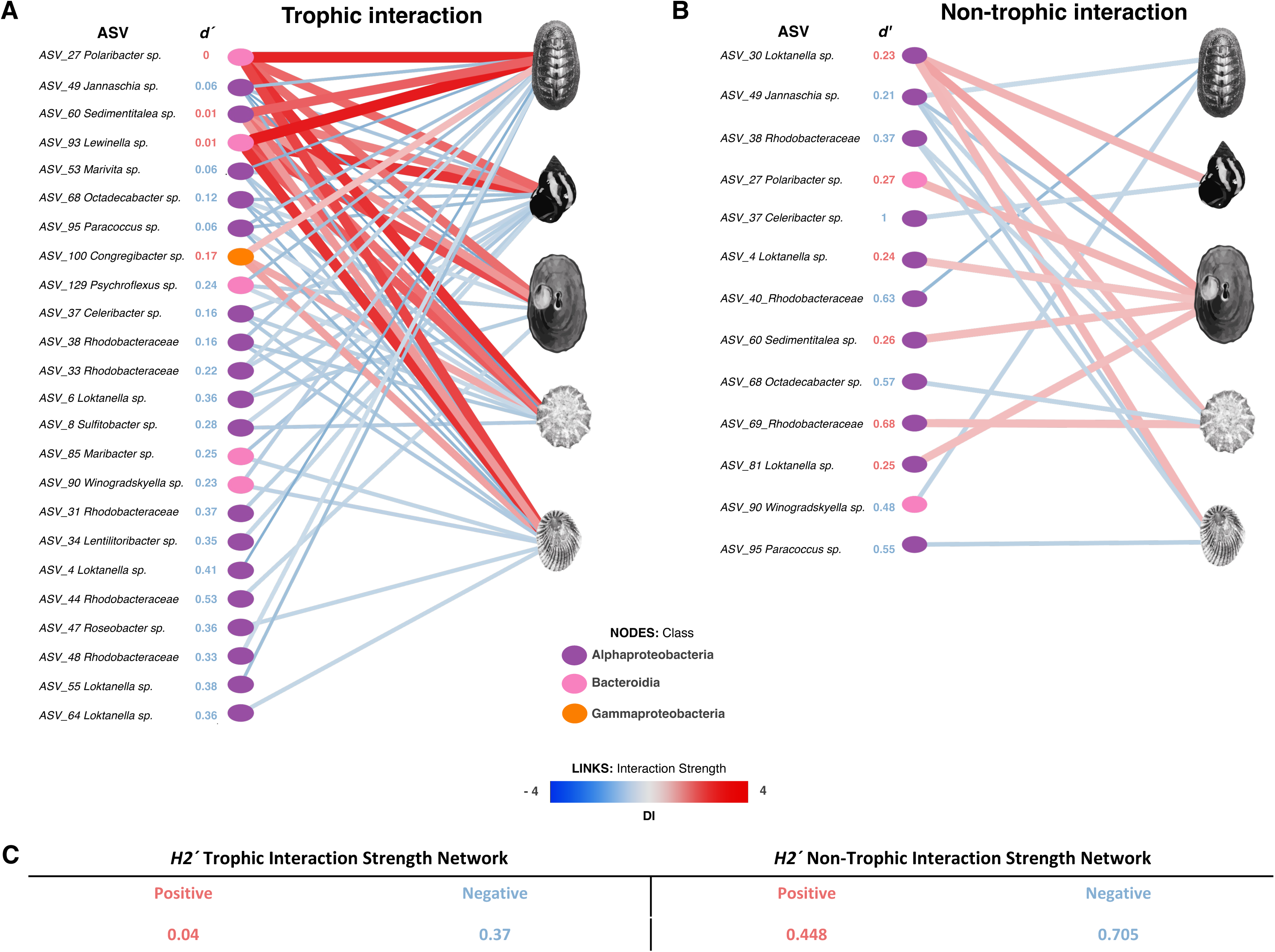
Bipartite network depicting interactions between molluscan grazer species and microbial groups (ASVs). The color and size of the links indicate the signs and values, respectively, of the interaction strength (per capita effect calculated by the Dynamic Index (DI)): --positive effect and --negative effect. A) Trophic interactions and B) non-trophic interactions. The specialization index (*d′*) of different bacterial groups within each positive or negative interaction strength during trophic or non-trophic interactions is displayed. C) Network-level specialization indices (*H_2_′*). Images credits: Ana María Valencia.

Within the Rhizobiacea Family (ASV_34 *Lentilitoribacter* sp.), only *E. peruviana* exhibited a negative per capita effect. In the Bacteroidia Class, the Sapropsiraceae Family (ASV_93 *Lewinella* sp.) displayed a positive per capita effect by all five grazer species. Conversely, the Flavobacteriaceae Family exhibited a positive per capita effect of ASV_27 (*Polaribacter* sp.) across all grazer species but a negative per capita effect on three ASVs (*Maribacter* sp.*, Psychroflexus* sp., and *Winogradskyella* sp.) specific to different grazers. Additionally, ASV_100 (*Heliaceaea* sp.) from Gammaproteobacteria showed a positive per capita effect by *C. granosus, S. araucana,* and *S. lessonii* (**Fig. 4a** and **Appendix S1: Fig. S5a**).

In contrast, the Flavobactriaceae Family exhibited a positive per capita effect in ASV_27 (*Polaribacter* sp.) across all five grazer species while displaying a negative per capita effect on three other ASVs (*Maribacter* sp.*, Psychroflexus* sp., and *Winogradskyella* sp.) in various grazers. Additionally, ASV_100 (*Heliaceaea* sp.) from the Gammaproteobacteria Class demonstrated a positive per capita effect among *C. granosus, S. araucana,* and *S.lessonii* (**Fig. 4a** and **Appendix S1: Fig. S5a**).

Following non-trophic interactions, the number of ASVs exhibiting a per capita effect by different grazers decreased compared to trophic interactions, with only 13 ASVs displaying positive or negative grazer effects (**Fig. 4b** and **Appendix S1: Fig. S5b**). The Rhodobacteraceae Family exhibited negative effects of grazers on six ASVs (*Celeribacter* sp.*, Jannaschia* sp.*, Octadecabacter* sp.*, Paracoccus* sp., and one unidentified ASV) with varying intensity. Within the Flavobacteriaceae Family, ASV_27 (*Polaribacter* sp.) demonstrated a per capita positive effect by *F. crassa*, while ASV_90 (*Winogradskyella* sp.) displayed a per capita negative effect by *C. granosus* (**Fig. 4b** and **Appendix S1: Fig. S5b**).

### Bipartite networks analyses

Upon constructing bipartite networks to analyze positive and negative interaction strengths between macrograzers and microbial ASVs using the Dynamic Index (DI), we evaluated the specialization indices of bipartite networks at the bacterial group level (*d′*) (**Fig. 4**). In the trophic positive interaction network, we observed that most ASVs displayed a generalized pattern (*d′* < 0.17). This was due to positive interactions between *Polaribacter* sp., *Sedimentitalea* sp., *Lewinella* sp., and all five macrograzers, as well as *Congregibacter* sp. interacting with three macrograzers. Conversely, the trophic negative interaction network exhibited a wide range of specialization, ranging from 0 (most generalized) to 0.53 (most specialized). Among the most generalized ASVs (*d′* < 0.12) were *Jannaschia* sp., *Marivita* sp., *Paracoccus* sp., and *Octadecabacter* sp. The most specialized ASVs (*d′* > 0.41) were ASV_44 from the Rhodobacteraceae family, and ASV_4 from *Loktanella* sp. (**Fig. 4a**).

In contrast, within the non-trophic positive interaction network, we observed a higher degree of specialization among ASVs. ASV_69 from the Rhodobacteraceae family exhibited a specialization index *d′* = 0.68, indicating a more specialized interaction pattern. Other ASVs displayed a relatively generalized pattern (*d′* ~ 0.2), including ASV_30, ASV_4, ASV_81 from *Loktanella* sp., *Sedimentitalea* sp., and *Polaribacter* sp. Moving to the non-trophic negative interaction network, we identified higher specialization values for certain ASVs. *Celeribacter* sp. and ASV_40 from the Rhodobacteraceae family displayed specialization indices of 1 and 0.63, respectively, indicating highly specialized interactions. On the other hand, ASV_38 from the Rhodobacteraceae family and *Jannaschia* sp. exhibited lower *d′* values (<0.37) and were considered more generalized, as they interacted with two or more macrograzers (**Fig. 4b**). The network-level specialization indices (*H_2_′*) (**Fig. 4c**) indicated lower values (*H_2_′* < 0.37) for the generalized trophic positive and negative interaction networks, while higher values (*H_2_′* > 0.45) were observed for the more specialized non-trophic positive and negative interaction networks. Higher values of *H_2_*, indicate increased nestedness, suggesting that species with fewer interactions tend to interact with subsets of species that have a higher number of interactions.

## DISCUSSION

Previous studies classified mollusk grazers into different functional groups due to their impact on algae (Lubchenco and Gaines 1981, Steneck and Dethier 1994, Duffy et al. 2001, Aguilera and Navarrete 2012) and microalgae (e.g., Bacillariophyta and Cyanobacteria) (Nicotri 1977, Jenkins et al. 2001, Aguilera et al. 2013). The most commonly used methodology for analyzing grazer effects on epilithic biofilms (periphyton) is measuring Chlorophyll-a content as an indicator of photosynthetic group biomass (Underwood 1984, Williams et al. 2000, Jenkins et al. 2001). However, some studies that compared periphyton Chlorophyll-a content with grazer density at different spatial and temporal scales did not show a relationship between grazing effort and microalgal abundance (Underwood 1984, Jenkins et al. 2001, Christofoletti et al. 2011). In contrast, experiments involving grazing and herbivore exclusion have demonstrated a negative effect on periphyton Chlorophyll-a content following trophic interactions with the entire grazer community (Williams et al. 2000), chitons over 25 days of grazing (Aguilera et al. 2013), littorinids over 40 days of grazing (Hidalgo et al. 2008), and limpets over one or two months of grazing (Valdivia et al. 2019). However, a positive effect on periphyton Chlorophyll-a content was observed in scurrinid and pulmonate limpets over 25 days of grazing (Aguilera et al. 2013). Our study found that after 24 hours, the Chlorophyll-a content methodology had limitations and only showed a negative per capita effect for one species, *F. crassa*.

In contrast, when using the epilithic biofilm cover methodology, we observed differential effects across all grazer treatments even within a short experiment duration. Another study utilizing image analysis techniques to assess biofilm cover on fiberglass panels found that optical density measurements were positively correlated with the total percentage cover of the biofilm (Anderson 1995). This suggests that the optical density method is an effective approach following two and four months of grazing by periwinkles, littorinids, and limpets. In our study, we introduce a novel protocol for analyzing the cover of epilithic biofilm. This protocol utilizes natural intertidal rocks as the substrate, ensuring a more realistic and reproducible assessment (Protocols.io: dx.doi.org/10.17504/protocols.io.e6nvw5epdvmk/v1) (Arboleda-Baena et al. 2023b).

Moreover, microscopy techniques have been employed in periphyton studies. For instance, bright field microscopy has been utilized for counting and identifying Bacillariophyta and Cyanobacteria groups (Aguilera et al. 2013), while Scanning Electron Microscopy (SEM) has been used to qualitatively assess differences between grazed and non-grazed areas (Nicotri 1977, Williams et al. 2000). In our study, SEM analysis revealed how the two grazers with the highest impact on epilithic biofilm cover exhibited contrasting grazing patterns due to their radula structure. *C. granosus*, with a Docoglossa radula, destroyed all remaining cells, whereas *F. crassa*, with a Steroglossa radula, left the remaining cells intact, potentially allowing for recolonization of the grazed area. Further studies employing microscopy should be conducted to assess the recolonization of grazed areas.

Previous studies have employed molecular tools to analyze the gut content of epilithic biofilms in a single species of littorinid and limpet grazers (Ding et al. 2018). However, these studies focused exclusively on oxygenic photoautotrophic groups. In contrast, our research enhances the resolution of bacterial communities and categorizes them based on their positive or negative interactions following 24 hours of grazing. Our approach revealed that not only do grazers with a higher effect on cover and Chlorophyll-a concentration exert a proportional impact on the most abundant microorganisms, but surprisingly, even grazers with minimal effects on cover and Chlorophyll-a can also demonstrate a significant impact on the abundance of ASVs within the biofilm. By examining the per capita grazing effects of the five most common species found in intertidal rocky shores, we constructed a high-resolution bipartite network. This network allowed us to identify non-interacting nodes and measure the intensity of interactions between two trophic levels. These results are crucial for distinguishing the strength of interactions among specific microbial taxonomic units and gaining a better understanding of the connections between the macroscopic and microscopic world. Our results contribute to enhancing our understanding of the complex interactions between microorganisms and macroorganisms in marine systems, particularly in relation to biofouling (Callow and Callow 2011, Navarrete et al. 2019) and the effective control of such processes (Navarrete et al. 2020, Arboleda-Baena et al. 2023a).

Previous studies have compared bipartite networks in various ecological contexts, including bacteria-microeukaryotes interactions (Fuhrman et al. 2015, Zheng et al. 2023), bacteria-virus interactions (Weitz et al. 2013, Fuhrman et al. 2015), symbiont-host, parasite-host, and/or predator-prey interactions in microorganisms (Bjorbækmo et al. 2020), as well as unicellular fungi-microorganisms (Moll et al. 2021), sponge-microorganisms (Thomas et al. 2016), plant-microorganism (Guo et al. 2009, Feng et al. 2019, Wang et al. 2023), and insect-microorganisms (Pechal and Benbow 2016) interactions. However, most of these networks have been constructed using correlation-based inference methods or literature, without analyzing the strength of interactions. What sets our bipartite network apart is in experimentally demonstrating how the interaction strength between grazers and microbial taxa (Amplicon Sequence Variants) varies and distinguishes between trophic and non-trophic interactions. This differentiation allows us to observe the differential impact of grazer guilds on microscopic communities, presenting a valuable insight into the dynamics of these interactions.

The application of specialized indices in bipartite networks(Blüthgen et al. 2006, 2007) revealed intriguing patterns of specialization at both the bacterial group level (d’) and the network level (H2’). Specifically, the non-trophic networks exhibited higher specialization, whereas the trophic networks displayed a more generalized nature. This finding suggests that non-trophic interactions may be directly influenced by the microbiota composition and chemical characteristics of the grazer’s pedal mucus. Supporting evidence for this hypothesis comes from previous research, which demonstrated significant variations in microbiota and pedal mucus chemistry among the five macrograzer species within the same locality (Arboleda-Baena et al. 2022). This variation likely contributes to the observed differences in specialization within the bipartite networks, underscoring the importance of these factors in shaping the dynamics of non-trophic interactions. Consequently, these driving factors may contribute to increased specialization of ASV indices (*d′*) within each grazer species. However, in the case of trophic interactions, we observed a higher degree of generalization in the ASVs associated with each grazer species. The main drivers of microbial compositional changes and the level of specialization in ASVs within trophic interactions were found to be the type of radula (feeding apparatus) and body size of the grazers. Interestingly, we did not detect any statistically significant differences in the specialization index at the bacterial group level (*d′*) between grazers after trophic interactions. This suggests that although the composition of ASVs may be influenced by specific grazer species, there is not a high level of specialization between the two trophic levels. These findings highlight the complex interplay between grazers, their feeding characteristics, body size and microbial communities, shedding light on the dynamics of trophic interactions in these systems.

Our results contribute towards unraveling the intricate network of interactions between macroorganisms and microorganisms in intertidal rocky shores, allowing us to capture the dynamics of this complex ecological system. Our findings provide compelling evidence of trophic interactions (consumptive effects) between grazers and epilithic biofilms, showcasing the substantial influence of these interactions on bacterial communities compared to non-trophic interactions (pedal mucus effects). Remarkably, our results reveal a clear pattern where trophic interactions predominantly result in positive effects on microbial abundance. This finding underscores the significant role that trophic interactions play in driving the dynamics and structure of microbial populations within this ecosystem. By shedding light on these relationships, our study advances our understanding of the interconnectedness and functioning of ecological networks within intertidal rocky shores. The magnitude of both trophic and non-trophic interactions exhibited substantial variation across different grazers and microbial groups, emphasizing the intricate nature of interactions between grazers and epilithic biofilms in marine systems. Notably, our analysis of bipartite networks revealed a higher degree of specialization in non-trophic interactions compared to trophic interactions, owing to distinct drivers such as the chemistry or microbiota of the pedal mucus specific to each grazer species.

This research complements the ecological network proposed by Kéfi and colleagues in 2015, as it demonstrates that mollusk grazers not only interact trophically with epilithic biofilms, as they had proposed, but also do so non-trophically. Additionally, it establishes that for future studies, three drivers that affect the assembly of microbial communities (**Fig. 5**) must be considered. These drivers depend on the type of interaction; for trophic interactions, the drivers are the type of radula and body size, while for non-trophic interactions, the pedal mucus plays a role. For future studies, it is important to consider these drivers to better understand the dynamics between macro and microorganisms in intertidal systems. By recognizing the inherent complexity of natural systems, this study successfully demonstrates the feasibility of integrating microbes into ecological networks. It is crucial to emphasize the challenging and pioneering nature of experimentally quantifying interactions with microbes, as it represents a significant contribution. Consequently, this research provides valuable insights and potential methodologies for quantifying such interactions across diverse species or systems, further underscoring its importance. Moreover, it highlights the need for further research to deepen our understanding of these intricate interactions and their implications.

**Fig. 5.**
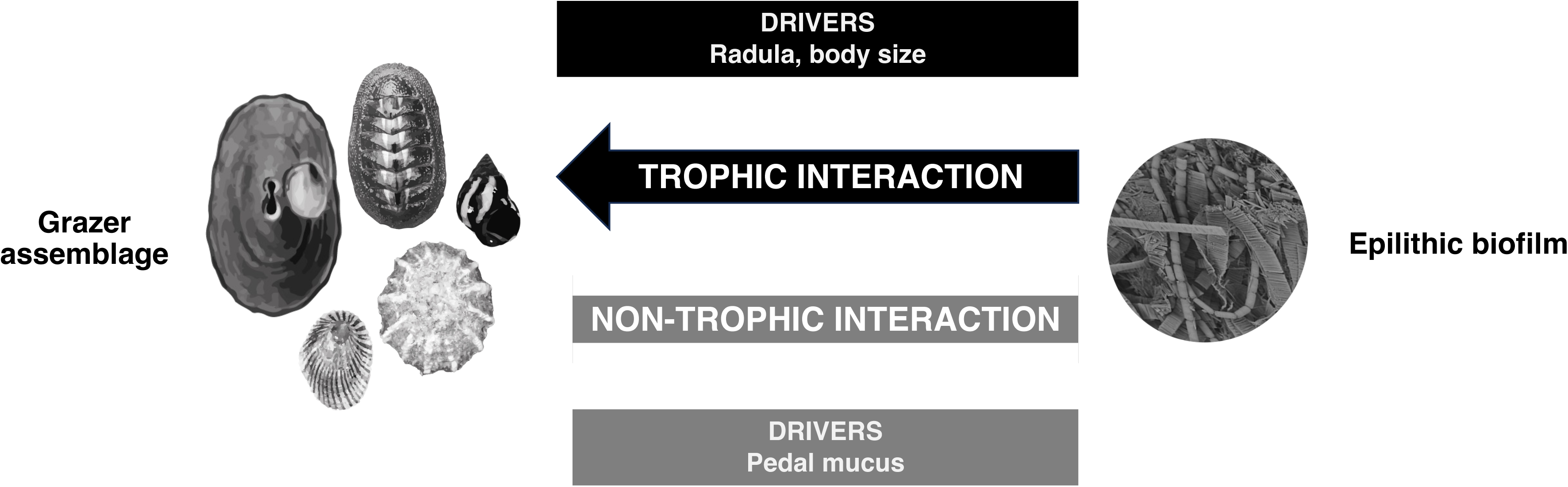
Conceptual model depicting the drivers of grazer assemblage that influence epilithic biofilm communities during trophic and non-trophic interactions in the intertidal rocky shore. Images/Photos credits: Ana María Valencia and Clara Arboleda-Baena.

## Acknowledgements

We thank Sergio Celis at ATIKA Ltda, Gadiel Alarcón at Season Mediciones Ambientales Limitada. We are also indebted to many students and research assistants at ECIM who collaborated with us in the field and laboratory. Funding for these studies and for international collaboration was provided by CONICYT (ANID) – National PhD scholarship Program 2016 to CMAB, and by Fondecyt grants No. 1160289 and 1200636 to SAN, and No. 1171259 to RDI. Complementary funding was provided by ANID PIA/BASAL FB0002 to SAN.

## Author Contribution

All authors agreed to be listed and have agreed on the submitted version of the manuscript. Conception and design of the work (CMAB, RDI, SAN), data collection (CMAB, BP, JP), data analysis and interpretation (CMAB, JP, HS, RL, RDI, SAN), drafting the article (CMAB, RDI, SAN), critical revision of the article (CMAB, EB, HD, RL, RDI, SAN), final approval of the version to be published (CMAB, BP, JP, EB, HS, RL, RD, SN).

## Conflict of Interest

The authors declare no conflict of interest.

## Notes

### Competing Interest Statement

The authors have declared no competing interest.

