## Supplementary Information for "Unifying microorganisms and macrograzers in rocky shore ecological networks"

**Appendix S1**

**Table S1.** Grazer body size of the five most abundant species in terms of total biomass: *Chiton granosus, Echinolittorina peruviana, Fissurella crassa, Scurria araucana*, and *Siphonaria lessonii*.

| SPECIES | N | Mean Body Size (mm) | sd | se | ci |
|---|---|---|---|---|---|
| ***C. granosus*** | 98 | 34.3 | 24.4 | 2.5 | 4.9 |
| ***E. peruviana*** | 400 | 10.1 | 3.3 | 0.2 | 0.3 |
| ***F. crassa*** | 55 | 42.9 | 28.9 | 3.9 | 7.8 |
| ***S. araucana*** | 90 | 16.1 | 7.7 | 0.8 | 1.6 |
| ***S. lessonii*** | 396 | 8.7 | 2.6 | 0.1 | 0.3 |

**Appendix S1**

**Figure S1.** Diagram of the experiment conducted to quantify the trophic (grazing effect) and non-trophic (grazer pedal mucus effect) interactions between five of the most abundant intertidal grazers and epilithic biofilms. Photos credits: Clara Arboleda-Baena.

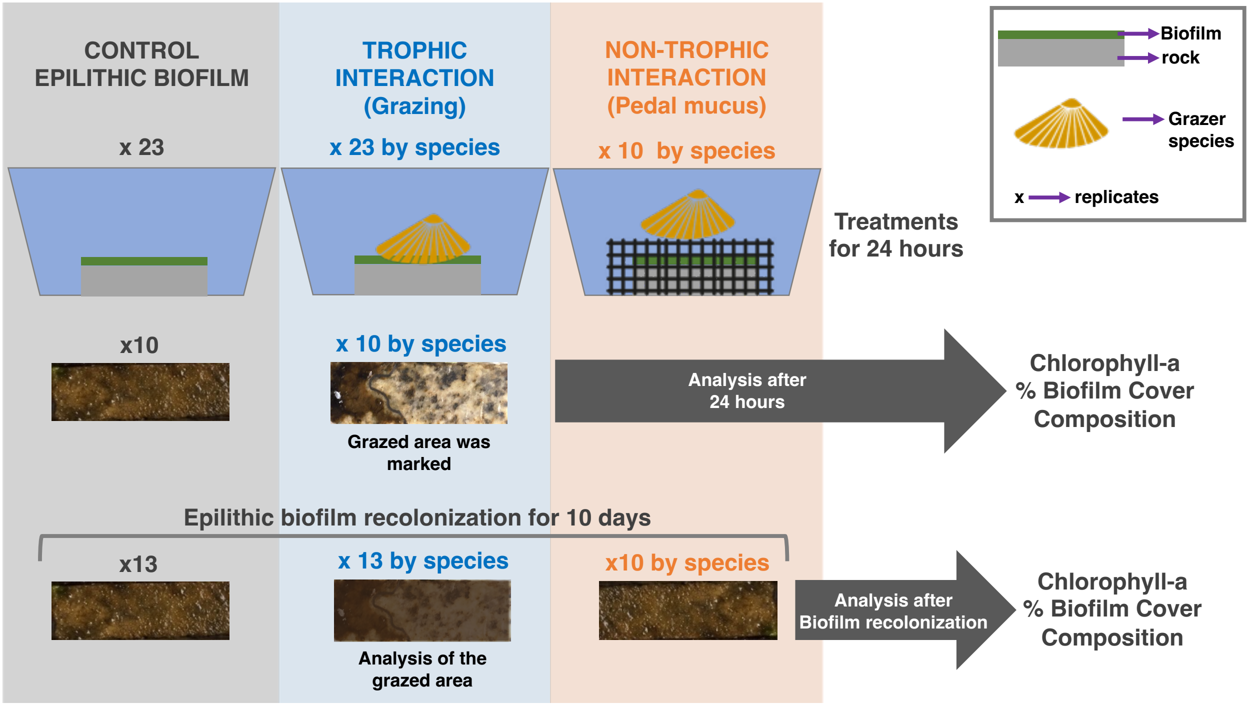


***Grazer effects on integrated measures of epilithic biofilm abundance***

Ten grazers (n = 10) of each species were chosen randomly and placed in individual clean aquaria (14.3 x 14.3 x 12.5 cm) with 400 mL filtered seawater to 0.22 µm (from the same location). Six grazing treatments (a-f) with 10 replicates each were implemented: a) Control: rock coupon with epilithic biofilm without grazers. Rock coupons with epilithic biofilms plus one individual of either b) *C. granosus*, c) *E. peruviana*, d) *F. crassa,* e) *S. araucana*, f) *S. lessonii*. Treatments were randomly assigned to the 60 experimental units (ten replicates per treatment). Every three hours, the temperature was checked, and feces were removed with sterile pipettes. The experiment lasted

24 hours, at which point the animals were carefully extracted. Within 24 hours, all 10 replicate rocks for each treatment were photographed, and the biofilm cover was analyzed using the software Image J with the protocol described in Protocols.io: https://dx.doi.org/10.17504/protocols.io.e6nvw5epdvmk/v1 (Arboleda-Baena et al. 2023). Biofilm total cover, expressed as a percentage of the rock coupon surface, provided an integrated estimate of grazers' total effect on autotroph and heterotroph biofilm components. Mean biofilm cover was compared among treatments with a Welch´s ANOVA (6 levels, fixed factor) due to deviations from homoscedasticity. Approximate normality was checked by visual inspection of residuals. To discern which treatment differed from others, a Games-Howell non-parametric post hoc test was used, which performs well under deviation of homoscedasticity (Ruxton and Beauchamp 2008). To assess effects only on autotrophs, and compare with previous studies on epilithic algae (Williams et al. 2000 p. 200, Aguilera et al. 2013, Valdivia et al. 2019), three replicate rocks of each treatment from the prior experiment were randomly selected for Chlorophyl-a analyses. Rock coupons were individually sonicated (Morris et al. 1998) to remove all biofilm which was recovered by filtration, preserved at 8ºC in 50 ml Falcon and sent to the SEASON Environmental Analysis Laboratory for quantification using standard Chlorophyll-*a* method (American Public Health Association. et al. 1998). Results expressed as Chlorophyll-*a* concentration from the entire rock coupon surface and comparison among treatments were conducted with Welch´s ANOVA (6 levels, fixed factor) after inspection for normality and detection of slightly heterogeneous variance. To observe and qualitatively assess grazing effects on the biofilm community we examined the grazed surface under an electron microscope. To this end, we chose one rock per treatment at random, cut a section of 1 x 1 x 1 cm with a diamond disk grinder, and immersed the piece for a few seconds in sterile seawater to remove the cut residue

that could affect the biofilm. The rock sample was then fixed in 4% glutaraldehyde buffered with sodium cacodylate (0.1 M, pH 7.2) at 4ºC. The coupon sections were scanned and photographed under a Hitachi TM3000 electron microscope at the Advanced Microscopy Laboratory (UMA) of the Pontificia Universidad Católica de Chile. To analyze the microbial composition of the epilithic biofilm at the experiment's onset, we obtained five replicates of the Control rock coupons (rock coupons with epilithic biofilm but without grazers). Each replicate was individually sonicated (Morris et al. 1998) to remove the biofilm, which was subsequently recovered by filtration using hydrophilic polyether sulfone filters with a pore size of 0.22 µm. The recovered biofilm was then preserved in liquid nitrogen at -196°C for subsequent DNA extraction and 16S rRNA-gene sequencing. A diagram of experiment is presented in **Appendix S1: Fig. S1.**

Aguilera, M., S. Navarrete, and B. Broitman. 2013. Differential effects of grazer species on periphyton of a temperate rocky shore. Marine Ecology Progress Series 484:63–78.

American Public Health Association., American Water Works Association., and Water Environment Federation. 1998. Standard methods for the examination of water and wastewater. APHA-AWWA-WEF, Washington, D.C.

Arboleda-Baena, C., C. B. Pareja, J. Poblete, E. Berlow, H. Sarmento, R. Logares, R. De la Iglesia, and S. Navarrete. 2023, July 13. Quantifying % Cover with ImageJ: An Analysis Tool for Image-based Assessments in Grazing Experiment Studies.

Morris, C. E., J.-M. Monier, and M.-A. Jacques. 1998. A Technique To Quantify the Population Size and Composition of the Biofilm Component in Communities of Bacteria in the Phyllosphere. Applied and Environmental Microbiology 64:4789–4795.

Ruxton, G. D., and G. Beauchamp. 2008. Time for some a priori thinking about post hoc testing. Behavioral Ecology 19:690–693.

Valdivia, N., L. M. Pardo, E. C. Macaya, P. Huovinen, and I. Gómez. 2019. Different ecological mechanisms lead to similar grazer controls on the functioning of periphyton Antarctic and sub-Antarctic communities. Progress in Oceanography 174:7–16.

Williams, G., M. Davies, and S. Nagarkar. 2000. Primary succession on a seasonal tropical rocky shore: the relative roles of spatial heterogeneity and herbivory. Marine Ecology Progress Series 203:81–94.

### Appendix S1

**Table S2.** Description, 16S rRNA sequencing characteristics of the samples.

| No. | ID | Grazer | Treatment | Number of QC reads | Number of ASVs | Number of reads after filtering | Number of ASVs after filtering |
|---|---|---|---|---|---|---|---|
| 1 | PositiveControl_72 | NA | New community Positive Control | 10633 | 174 | 6215 | 172 |
| 2 | PositiveControl_78 | NA | New community Positive Control | 13483 | 195 | 6215 | 193 |
| 3 | PositiveControl_79 | NA | New community Positive Control | 36465 | 376 | 6215 | 336 |
| 4 | PositiveControl_80 | NA | New community Positive Control | 24573 | 338 | 6215 | 311 |
| 5 | Cg_162M | *C. granosus* | Pedal Mucus Control | 12240 | 321 | 6215 | 312 |
| 6 | Cg_166M | *C. granosus* | Pedal Mucus Control | 14785 | 389 | 6215 | 365 |
| 7 | Cg_167M | *C. granosus* | Pedal Mucus Control | 8284 | 183 | 6215 | 182 |
| 8 | Cg_169M | *C. granosus* | Pedal Mucus Control | 16104 | 371 | 6215 | 343 |
| 9 | Ep_172M | *E. peruviana* | Pedal Mucus Control | 9455 | 287 | 6215 | 277 |
| 10 | Ep_173M | *E. peruviana* | Pedal Mucus Control | 7077 | 183 | 6215 | 183 |
| 11 | Ep_174M | *E. peruviana* | Pedal Mucus Control | 7838 | 203 | 6215 | 203 |
| 12 | Ep_176M | *E. peruviana* | Pedal Mucus Control | 8912 | 158 | 6215 | 157 |
| 13 | Ep_177M | *E. peruviana* | Pedal Mucus Control | 12961 | 212 | 6215 | 204 |
| 14 | Ep_179M | *E. peruviana* | Pedal Mucus Control | 7013 | 143 | 6215 | 143 |
| 15 | Ep_180M | *E. peruviana* | Pedal Mucus Control | 13637 | 233 | 6215 | 225 |
| 16 | Fcpg_141M | *F.crassa* | Pedal Mucus Control | 11518 | 221 | 6215 | 212 |
| 17 | Fcpg_146M | *F.crassa* | Pedal Mucus Control | 13764 | 238 | 6215 | 228 |
| 18 | Fcpg_148M | *F.crassa* | Pedal Mucus Control | 6215 | 197 | 6215 | 197 |
| 19 | Sa_151M | *S. araucana* | Pedal Mucus Control | 17449 | 267 | 6215 | 248 |
| 20 | Sa_152M | *S. araucana* | Pedal Mucus Control | 26433 | 285 | 6215 | 249 |
| 21 | Sa_153M | *S. araucana* | Pedal Mucus Control | 6663 | 141 | 6215 | 141 |
| 22 | Sa_157M | *S. araucana* | Pedal Mucus Control | 6820 | 141 | 6215 | 141 |
| 23 | Sa_159M | *S. araucana* | Pedal Mucus Control | 16041 | 226 | 6215 | 213 |
| 24 | Sl_182M | *S.lessonii* | Pedal Mucus Control | 16251 | 230 | 6215 | 214 |
| 25 | Sl_183M | *S.lessonii* | Pedal Mucus Control | 12183 | 242 | 6215 | 236 |
| 26 | Sl_184M | *S.lessonii* | Pedal Mucus Control | 13181 | 217 | 6215 | 203 |
| 27 | Sl_185M | *S.lessonii* | Pedal Mucus Control | 16622 | 225 | 6215 | 211 |
| 28 | Sl_186M | *S.lessonii* | Pedal Mucus Control | 23632 | 346 | 6215 | 305 |
| 29 | Sl_187M | *S.lessonii* | Pedal Mucus Control | 27171 | 356 | 6215 | 310 |
| 30 | Sl_188M | *S.lessonii* | Pedal Mucus Control | 14684 | 248 | 6215 | 237 |
| 31 | Sl_189M | *S.lessonii* | Pedal Mucus Control | 8234 | 205 | 6215 | 204 |
| 32 | Sl_190M | *S.lessonii* | Pedal Mucus Control | 7938 | 278 | 6215 | 277 |
| 33 | Cg_101 | *C. granosus* | New community of grazed rock | 38590 | 687 | 6215 | 561 |

| | | | | | | | |
|---|---|---|---|---|---|---|---|
| 34 | Cg_103 | *C. granosus* | New community of grazed rock | 28213 | 529 | 6215 | 449 |
| 35 | Cg_104 | *C. granosus* | New community of grazed rock | 16511 | 398 | 6215 | 380 |
| 36 | Cg_109 | *C. granosus* | New community of grazed rock | 8730 | 368 | 6215 | 364 |
| 37 | Cg_110 | *C. granosus* | New community of grazed rock | 24743 | 491 | 6215 | 423 |
| 38 | Ep_121 | *E. peruviana* | New community of grazed rock | 10210 | 263 | 6215 | 263 |
| 39 | Ep_122 | *E. peruviana* | New community of grazed rock | 28927 | 662 | 6215 | 565 |
| 40 | Ep_123 | *E. peruviana* | New community of grazed rock | 20056 | 508 | 6215 | 467 |
| 41 | Ep_124 | *E. peruviana* | New community of grazed rock | 26654 | 626 | 6215 | 542 |
| 42 | Ep_125 | *E. peruviana* | New community of grazed rock | 13408 | 398 | 6215 | 376 |
| 43 | Ep_127 | *E. peruviana* | New community of grazed rock | 9580 | 327 | 6215 | 325 |
| 44 | Ep_129 | *E. peruviana* | New community of grazed rock | 26492 | 682 | 6215 | 575 |
| 45 | Ep_130 | *E. peruviana* | New community of grazed rock | 31844 | 691 | 6215 | 566 |
| 46 | Fcg_82 | *F.crassa large* | New community of grazed rock | 11388 | 339 | 6215 | 332 |
| 47 | Fcg_83 | *F.crassa large* | New community of grazed rock | 15848 | 439 | 6215 | 416 |
| 48 | Fcg_87 | *F.crassa large* | New community of grazed rock | 28877 | 620 | 6215 | 516 |
| 49 | Fcg_89 | *F.crassa large* | New community of grazed rock | 22996 | 523 | 6215 | 457 |
| 50 | Fcp_113 | *F.crassa small* | New community of grazed rock | 15044 | 448 | 6215 | 424 |
| 51 | Fcp_114 | *F.crassa small* | New community of grazed rock | 18397 | 635 | 6215 | 565 |
| 52 | Fcp_120 | *F.crassa small* | New community of grazed rock | 10167 | 664 | 6215 | 646 |
| 53 | Sa_91 | *S. araucana* | New community of grazed rock | 13762 | 387 | 6215 | 367 |
| 54 | Sa_92 | *S. araucana* | New community of grazed rock | 21903 | 460 | 6215 | 420 |
| 55 | Sa_93 | *S. araucana* | New community of grazed rock | 11404 | 435 | 6215 | 425 |
| 56 | Sa_94 | *S. araucana* | New community of grazed rock | 12833 | 519 | 6215 | 487 |
| 57 | Sa_95 | *S. araucana* | New community of grazed rock | 23034 | 850 | 6215 | 776 |
| 58 | Sa_96 | *S. araucana* | New community of grazed rock | 22585 | 550 | 6215 | 497 |
| 59 | Sa_97 | *S. araucana* | New community of grazed rock | 7691 | 251 | 6215 | 249 |
| 60 | Sa_98 | *S. araucana* | New community of grazed rock | 15286 | 453 | 6215 | 430 |
| 61 | Sa_100 | *S. araucana* | New community of grazed rock | 17145 | 396 | 6215 | 377 |
| 62 | Sl_131 | *S.lessonii* | New community of grazed rock | 9359 | 344 | 6215 | 338 |
| 63 | Sl_132 | *S.lessonii* | New community of grazed rock | 19658 | 499 | 6215 | 457 |
| 64 | Sl_135 | *S.lessonii* | New community of grazed rock | 12593 | 329 | 6215 | 313 |
| 65 | Sl_136 | *S.lessonii* | New community of grazed rock | 9892 | 309 | 6215 | 298 |
| 66 | Sl_137 | *S.lessonii* | New community of grazed rock | 9940 | 427 | 6215 | 417 |
| 67 | Sl_138 | *S.lessonii* | New community of grazed rock | 10271 | 332 | 6215 | 328 |

**Appendix S1. 2**

**Figure S2.** Rarefaction curves. Sample size and ASVs are presented by treatments ■ Epilithic biofilm control, ■ Pedal mucus control, ■ Bacterial communities on grazed rock.

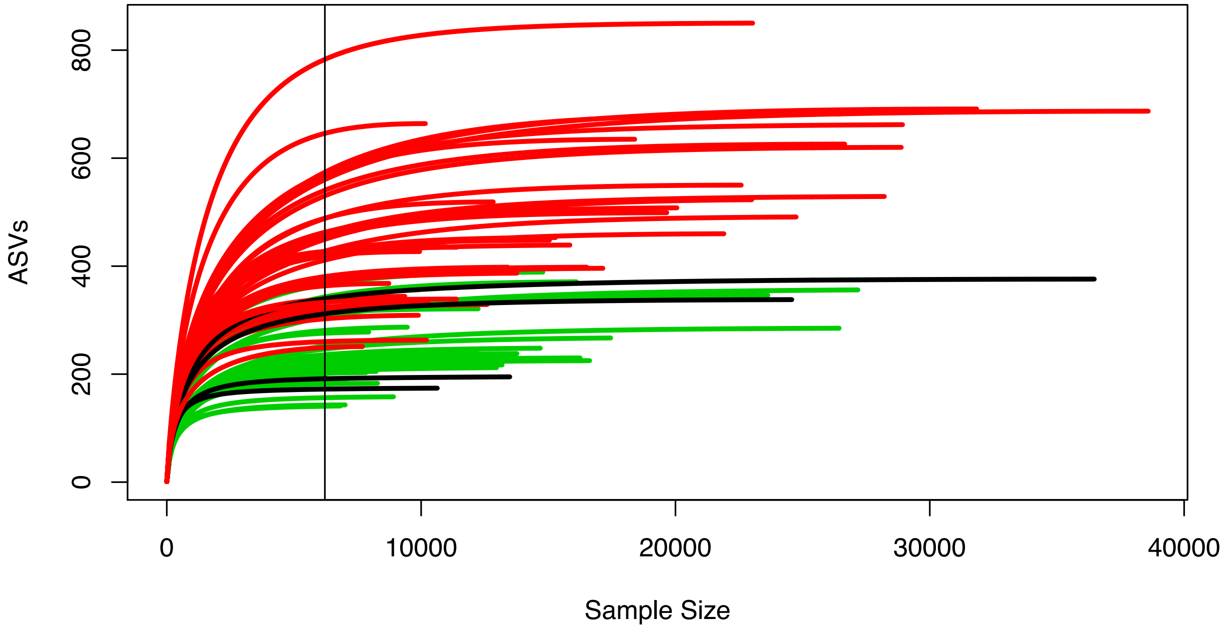

**Appendix S1**

**Figure S3.** Interaction strength analyses with Chlorophyll-*a* content and cover after A,C) 24 hours of grazing and B,D) 10 days of epilithic biofilm recolonization of the grazed area. Bars represent 95% confidence intervals estimated through bootstrapping procedure.

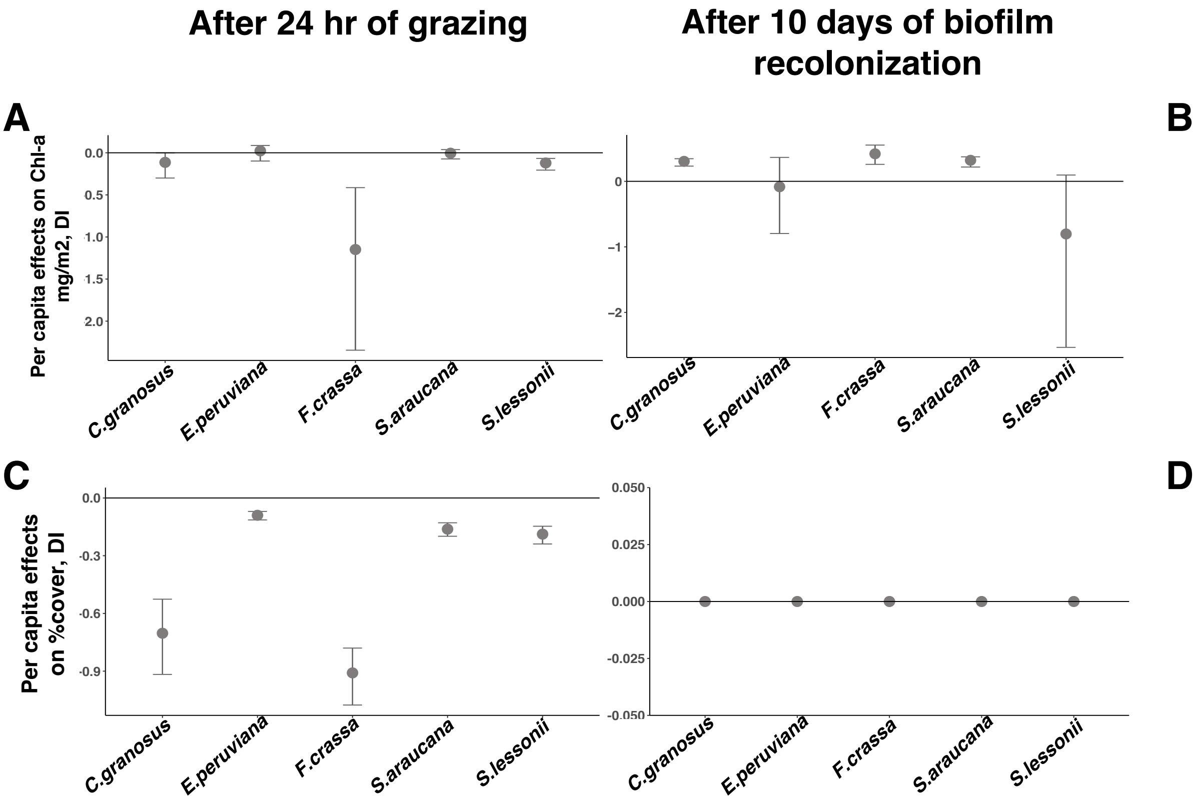

**Appendix S1**

**Table S3.** Grazing effect over cover biofilm percentage. Games-Howell post hoc tests at the experiment-wise error rate = 0.05.

| | diff | ci.lo | ci.hi | t | df | p |
|---|---|---|---|---|---|---|
| Control-*C. granosus* | 45.89 | 30.66 | 61.12 | 9.504 | 19.3 | <.001 |
| *E. peruviana-C. granosus* | 37.37 | 21.96 | 52.77 | 7.598 | 20.7 | <.001 |
| *F. crassa-C. granosus* | -9.77 | -26.62 | 7.08 | 1.756 | 32.3 | .507 |
| *S. araucana-C. granosus* | 31.22 | 15.54 | 46.89 | 6.182 | 22.9 | <.001 |
| *S. lessonii-C. granosus* | 29.18 | 13.21 | 45.15 | 5.631 | 25.0 | <.001 |
| *E. peruviana*-Control | -8.53 | -11.94 | -5.11 | 7.679 | 25.4 | <.001 |
| *F. crassa*-Control | -55.67 | -64.13 | -47.20 | 19.650 | 40.8 | <.001 |
| *S. araucana*-Control | -14.67 | -19.65 | -9.70 | 9.185 | 21.9 | <.001 |
| *S. lessonii*-Control | -16.71 | -22.89 | -10.53 | 8.460. | 20.9 | <.001 |
| *F. crassa-E. peruviana* | -47.14 | -55.99 | -38.29 | 15.809 | 48.4 | <.001 |
| *S. araucana-E. peruviana* | -6.15 | -11.74 | -0.56 | 3.326 | 33.1 | .024 |
| *S. lessonii-E. peruviana* | -8.19 | -14.84 | -1.53 | 3.749 | 28.9 | .009 |
| *S. araucana-F. crassa* | 40.99 | 31.56 | 50.42 | 12.828. | 55.7 | <.001 |
| *S. lessonii-F. crassa* | 38.95 | 28.93 | 48.97 | 11.457 | 58.0 | <.001 |
| *S. lessonii-S. araucana* | -2.04 | -9.46 | 5.38 | 0.826 | 36.2 | 0.961 |

**Appendix S1.**

**Figure S4.** Scanning electron microcopy after the grazing experiment of five most abundant species of the intertidal rocky shore. Treatments: 1) Control biofilm without grazing, 2) Control Rock surface, grazing of 3) *C.granosus,* 4) *E.peruviana,* 5) *F.crassa* 6) *S.araucana y* 7) *S.lessonii.* Photos credits: Clara Arboleda-Baena and Claudia Belén Pareja.

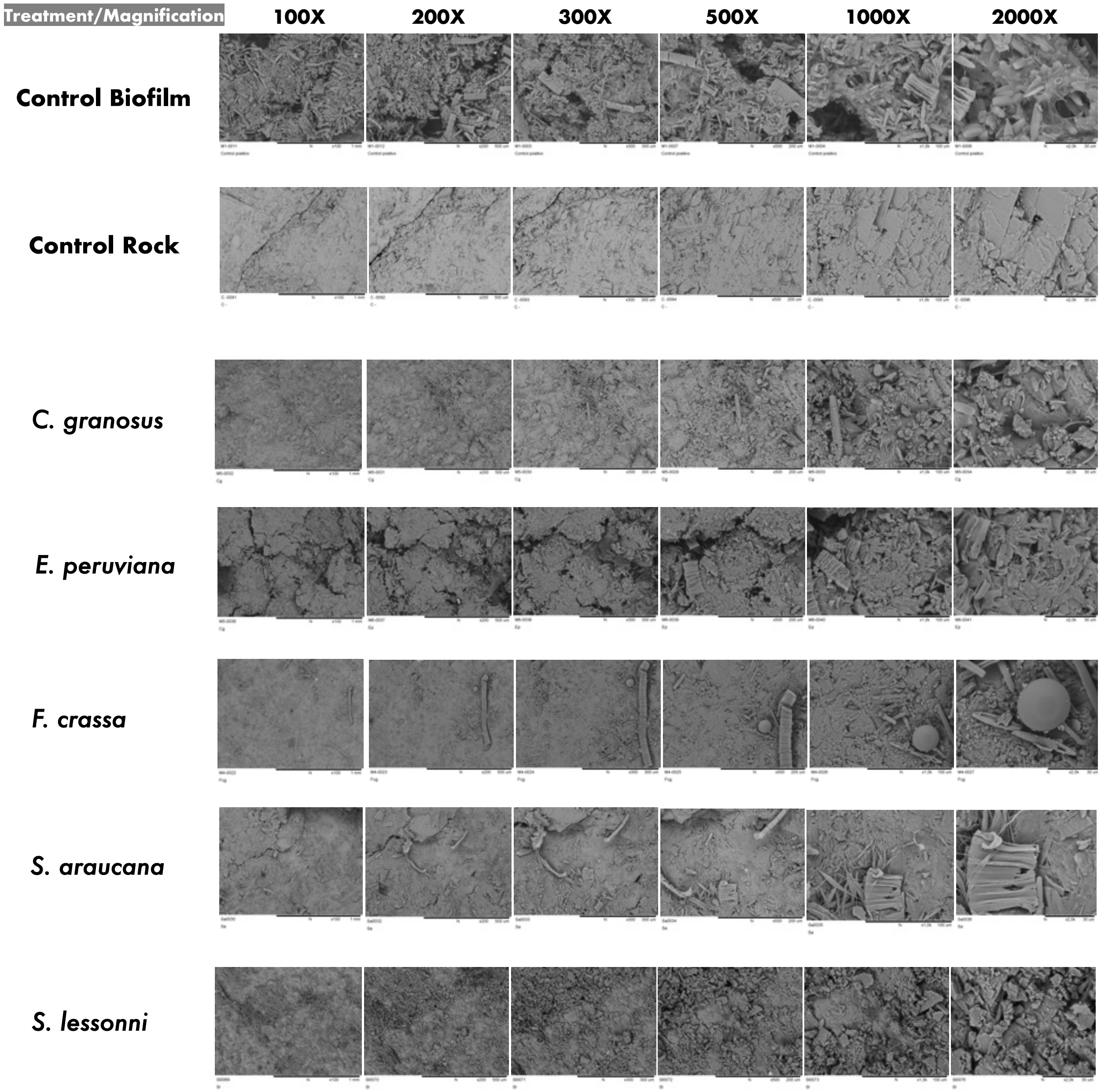

**Appendix S1.**

**Table S4.** False Discovery Rate (FDR) pairwise comparisons of bacterial communities after total, non-trophic and trophic interaction with *C. granosus, E. peruviana, F. crassa, S. araucana* and *S. lessonii*. The distances used were Bray-Curtis.

a. **NON-TROPHIC INTERACTION (NTI)**

| **PERMANOVA = 0.08092** |
|---|

b. **TROPHIC INTERACTION (TI)**

| **PERMANOVA = 0.02797*** | | | | | | | |
|---|---|---|---|---|---|---|---|
| | **pairs** | **Df** | **SumsOfSqs** | **F.Model** | **R2** | **p.value** | **p.adjusted** |
| **1** | *S. araucana vs C. granosus* | 1 | 0.300 | 1.257 | 0.095 | 0.144 | 0.274 |
| **2** | *S. araucana vs F.crassa* | 1 | 0.280 | 1.102 | 0.073 | 0.277 | 0.308 |
| **3** | *S. araucana vs E. peruviana* | 1 | 0.325 | 1.434 | 0.087 | 0.053 | 0.218 |
| **4** | *S. araucana vs S.lessoni* | 1 | 0.379 | 1.518 | 0.105 | 0.019 | 0.190 |
| **5** | *C. granosus vs F.crassa* | 1 | 0.309 | 1.193 | 0.107 | 0.185 | 0.274 |
| **6** | *C. granosus vs E. peruviana* | 1 | 0.219 | 0.990 | 0.083 | 0.418 | 0.418 |
| **7** | *C. granosus vs S.lessoni* | 1 | 0.291 | 1.151 | 0.113 | 0.219 | 0.274 |
| **8** | *F.crassa vs E. peruviana* | 1 | 0.317 | 1.318 | 0.092 | 0.087 | 0.218 |
| **9** | *F.crassa vs S.lessoni* | 1 | 0.344 | 1.277 | 0.104 | 0.066 | 0.218 |
| **10** | *E. peruviana vs S.lessoni* | 1 | 0.266 | 1.136 | 0.086 | 0.204 | 0.274 |

**Appendix S1.**

**Table S5.** ANOVA, KRUSKAL-WALLIS test and Tukey post hoc test multiple comparisons of bacterial communities' richness and diversity after total, non-trophic and trophic interaction with *C. granosus, E. peruviana, F. crassa, S. araucana* and *S. lessonii*. 95% family-wise confidence level.

**a. NON-TROPHIC INTERACTION (NTI) Richness**

ANOVA

| | Df. | Sum Sq | Mean Sq | F value | Pr(>F) |
|---|---|---|---|---|---|
| Information | 4 | 24701 | 6175 | 3.156 | 0.0331 * |
| Residuals | 23 | 45000 | 1957 | | |

TUKEY POST HOC TEST

| | diff | lwr | upr | p adj |
|---|---|---|---|---|
| *E. peruviana-C. granosus* | -83.214286 | -165.16809 | -1.2604794 | 0.0453792 |
| *F. crassa-C. granosus* | -71.166667 | -171.03095 | 28.6976189 | 0.2511927 |
| *S. araucana-C. granosus* | -87.900000 | -175.61181 | -0.1881928 | 0.0493307 |
| *S. lessonii-C. granosus* | -43.833333 | -122.40613 | 34.7394664 | 0.4830495 |
| *F. crassa-E. peruviana* | 12.047619 | -78.18055 | 102.2757903 | 0.9945074 |
| *S. araucana-E. peruviana* | -4.685714 | -81.24686 | 71.8754278 | 0.9997409 |
| *S. lessonii-E. peruviana* | 39.380952 | -26.51239 | 105.2742920 | 0.4157901 |
| *S. araucana-F. crassa* | -16.733333 | -112.22185 | 78.7551876 | 0.9846527 |
| *S. lessonii-F. crassa* | 27.333333 | -59.83536 | 114.5020281 | 0.8835266 |
| *S. lessonii-S. araucana* | 44.066667 | -28.86390 | 116.9972292 | 0.4050864 |

**b. NON-TROPHIC INTERACTION (NTI) Diversity**

ANOVA

| | Df | Sum Sq | Mean Sq | F valu | Pr(>F) |
|---|---|---|---|---|---|
| Information | 4 | 0.8119 | 0.20297 | 2.403 | 0.0792 . |
| Residuals | 23 | 1.9430 | 0.08448 | | |

**c. TROPHIC INTERACTION (TI) Richness**

KRUSKAL-WALLIS.
Kruskal-Wallis rank sum test
data: Richness by Information
Kruskal-Wallis chi-squared = 4.9898, *df* = 4, p-value = 0.2883

**d. TROPHIC INTERACTION (TI) Diversity**

ANOVA

| | Df | Sum Sq | Mean Sq. | F value | Pr(>F) |
|---|---|---|---|---|---|
| Information | 4 | 0.1894 | 0.04736 | 0.574 | 0.684 |
| Residuals | 30 | 2.4755 | 0.08252 | | |

**Appendix S1.**

**Figure S5.** Interaction strength (per capita effect calculated by the Dynamic Index (DI)) between macrograzers and the most abundant microbial groups (Total number of reads > 0.5%). *Per capita* effect of *C.* granosus, *E. peruviana, F. crassa, S. araucana* and *S. lessonii* during the A) Trophic interaction (grazing consumption), B) Non-trophic interaction (pedal mucus effect). Bars indicated the 95% confidence intervals estimated through bootstrapping procedure. Class names in different colors and Family names in parentheses.

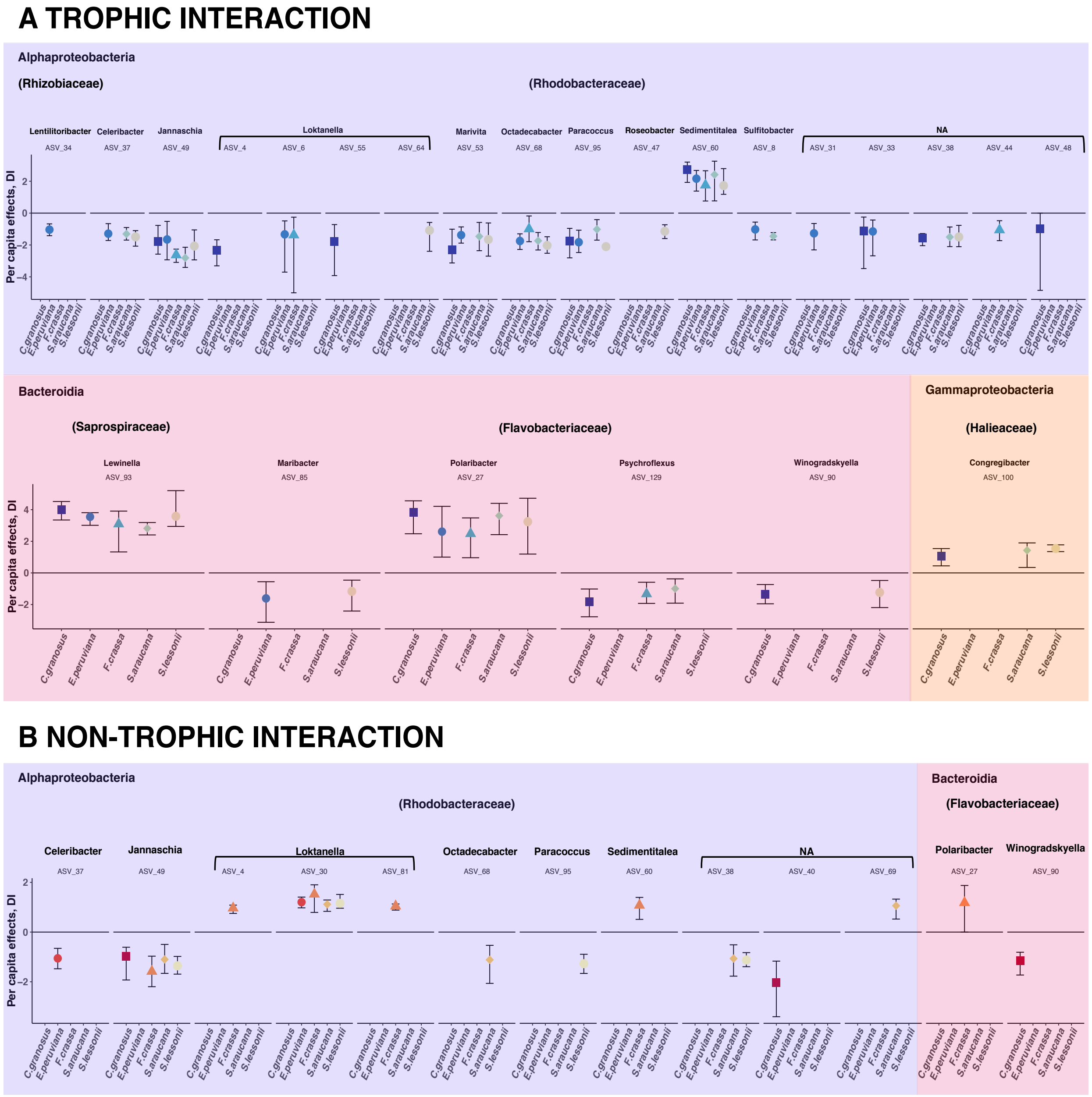
